# Phages reconstitute NAD^+^ to counter bacterial immunity

**DOI:** 10.1101/2024.02.11.579819

**Authors:** Ilya Osterman, Hadar Samra, Francois Rousset, Elena Loseva, Maxim Itkin, Sergey Malitsky, Erez Yirmiya, Adi Millman, Rotem Sorek

## Abstract

Bacteria defend against phage infection via a variety of antiphage defense systems. Many defense systems were recently shown to deplete cellular nicotinamide adenine dinucleotide (NAD^+^) in response to infection, by breaking NAD^+^ to ADP-ribose (ADPR) and nicotinamide. It was demonstrated that NAD^+^ depletion during infection deprives the phage from this essential molecule and impedes phage replication. Here we show that a substantial fraction of phages possess enzymatic pathways allowing reconstitution of NAD^+^ from its degradation products in infected cells. We describe NAD^+^ reconstitution pathway 1 (NARP1), a two-step pathway in which one enzyme phosphorylates ADPR to generate ADPR-pyrophosphate (ADPR-PP), and the second enzyme conjugates ADPR-PP and nicotinamide to generate NAD^+^. Phages encoding the NARP1 pathway can overcome a diverse set of defense systems, including Thoeris, DSR1, DSR2, SIR2-HerA, and SEFIR, all of which deplete NAD^+^ as part of their defensive mechanism. Phylogenetic analyses show that NARP1 is primarily encoded on phage genomes, suggesting a phage-specific function in countering bacterial defenses. A second pathway, NARP2, allows phages to overcome bacterial defenses by building NAD^+^ via metabolites different than ADPR-PP. Our findings report a unique immune evasion strategy where viruses rebuild molecules depleted by defense systems, thus overcoming host immunity.

## Introduction

Nicotinamide adenine dinucleotide (NAD^+^/NADH) is a central metabolite essential for numerous core metabolic processes across all domains of life. In most bacteria NAD^+^ is an essential cofactor for redox reactions^1^, and in the absence of NAD^+^ pathways such as oxidative phosphorylation^2^, amino acids biosynthesis^3^, and fatty acids biosynthesis^3^, are arrested. NAD^+^ was also shown to be necessary for post-translational protein modifications^4^ and for DNA ligation processes^5^.

Recent data show that NAD^+^ metabolism is central for bacterial defense against phages^6-9^. Specifically, numerous bacterial defense systems were shown to deplete cellular NAD^+^ once they detect phage infection, thus depleting the cell of energy and impeding phage propagation. Such defense systems include prokaryotic argonautes (pAgo)^6,9^, type I Thoeris^7^, AVAST^10^, CBASS^11^, DSR1^6^, DSR2^6^, SIR2-HerA^6^, SEFIR^8^, and additional defense systems encoding protein domains associated with NAD^+^ depletion^12^. A recent analysis of the abundance of defense systems in microbial genomes show that at least 7% of all sequenced bacterial genomes carry defense systems that cause NAD^+^ depletion in response to phage infection^8^.

Once NAD^+^-depleting defense systems detect phage infection, they break down NAD^+^ to ADP-ribose (ADPR) and nicotinamide^6^. As there is no known cellular pathway that can directly re-build NAD^+^ from these molecules, breaking of NAD^+^ into ADPR and nicotinamide is an efficient way to deplete NAD^+^ from infected cells. Depletion of NAD^+^ during infection was shown to halt phage propagation and, in some cases, cause premature cell lysis^6,7^, possibly by activating the lysis machinery of the phage prior to the completion of the phage cycle^6,13^.

Here we show that at least 5% of sequenced phage genomes encode NAD^+^ reconstitution pathways that allow them to rebuild NAD^+^ directly from ADPR and nicotinamide. We discover NAD^+^ reconstitution pathway 1 (NARP1), a phage-encoded pathway involving two enzymatic reactions that were not described before. We also show that NARP2, an alternative NAD^+^ reconstitution pathway utilizing classical NAD^+^-salvage enzymatic reactions, is used by phages to rebuild NAD^+^ during infection. NAD^+^ reconstitution pathways allow phages to overcome multiple NAD^+^-depleting defense systems irrespective of the mechanism of phage detection and signal transfer in these systems.

## Results

### A two-gene operon protects phages from NAD^+^-depleting bacterial defenses

While examining the genomes of phages from the BASEL collection^14^ we noticed a two-gene operon that recurred in four phages in this set (Figure 1A). This operon caught our attention because the pfam annotations of its two genes suggested involvement in synthesis of biochemical intermediates in the NAD^+^ biosynthesis pathway. The first gene in this operon was annotated as belonging to the **p**hospho**r**ibosylpyrophosphate **s**ynthetase family, Prs (pfam accession PF14572), and the second gene was annotated as **n**icotin**am**ide **p**hosphoribosyl**t**ransferase, Nampt (pfam accessions PF18127 and PF04095) (Figure 1A). In bacteria, Prs enzymes produce **p**hospho**r**ibosyl**p**yro**p**hosphate (PRPP), a biochemical intermediate in the biosynthesis pathway of purine and pyrimidine nucleotides, as well as NAD^+^. Nampt enzymes utilize PRPP as a substrate and react it with nicotinamide to generate **n**icotinamide **m**ono**n**ucleotide (NMN), a molecular moiety that can be used as a precursor for NAD^+^ biosynthesis via NAD^+^ salvage pathways (Figure 1B, 1C). It was previously shown that genes annotated as nicotinamide phosphoribosyltransferases are abundant in phage genomes^15^.

**Figure 1.**
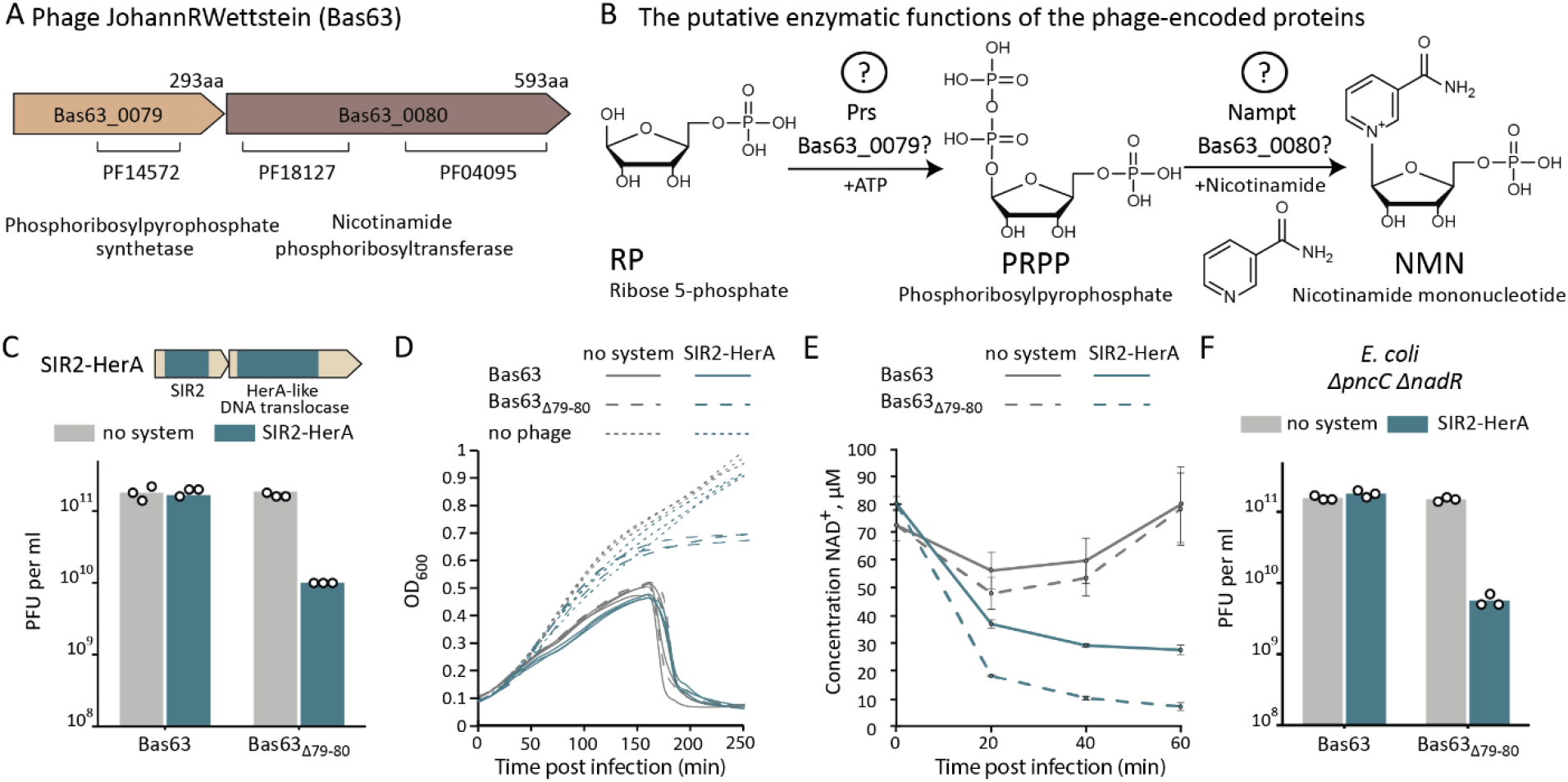
A phage-encoded operon that counteracts bacterial NAD^+^ depletion defense. **A**. Domain organization of genes 79-80 from phage JohannRWettstein (Bas63). Pfam annotations are indicated. **B**. The putative enzymatic functions of the phage-encoded proteins according to their predicted protein domains. Later in the paper we show that these predicted reactions are not the actual reactions performed by the phage enzymes. **C**. Operon 79-80 allows phage Bas63 to overcome SIR2-HerA defense. Data represent plaque-forming units per millilitre (PFU per ml) of wild type Bas63 and Bas63 in which the two-gene operon was deleted, infecting cells that express the SIR2-HerA defense system or control cells with an empty vector (no system). Bar graph represents average of three independent replicates, with individual data points overlaid. **D**. Liquid culture growth of *E. coli* cells expressing the SIR2-HerA operon or control strain without defense genes (no system). Cells were infected by phages Bas63 and Bas63_Δ79-80_ at MOI=0.01. Data from three replicates are presented as individual curves. OD_600_, optical density at 600 nm. **E**. NAD^+^ concentration in the lysate of cells expressing SIR2-HerA or control cells with an empty vector instead (no system), infected with phages Bas63 and Bas63_Δ79-80_ at MOI=10. Cells were collected before infection (0 min) and 20, 40 or 60 minutes after infection. NAD^+^ concentration in cell lysates was measured by the NAD/NADH-Glo biochemical assay. The experiment was performed in three replicates, error bars represent standard deviation. **F**. Same experiment as in panel D, but made on an *E. coli* strain with deletions of the two genes *pncC* and *nadR*.

We hypothesized that phages utilize this two-gene operon to overcome the effects of NAD^+^-depleting defense systems. To test this hypothesis, we deleted the two-gene operon from the BASEL phage Bas63 (JohannRWettstein^14^). The wild type Bas63 was able to infect *E. coli* cells encoding the defense system SIR2-HerA (also called Nezha^16^), which is known to deplete NAD^+^ following phage recognition^6,16^ (Figure 1C). In contrast, a Bas63 strain in which the two-gene operon was deleted formed ∼20-fold less plaques on agar plates, suggesting that SIR2-HerA defends against the modified phage and that the phage two-gene operon can counter this defense (Figure 1C). These results were confirmed by infecting cells in liquid culture, where a culture of bacteria encoding the SIR2-HerA defense system collapsed when infected with the WT phage, but was able to grow when infected by the phage in which the two genes were deleted (Figure 1D).

To test if the two-gene operon allows the phage to overcome the NAD^+^ depletion effects of SIR2-HerA, we recorded cellular NAD^+^ levels during infection. NAD^+^ was severely depleted in cells expressing SIR2-HerA that were infected by the mutated phage, confirming the activity of SIR2-HerA against this phage (Figure 1E). When infected with the wild type Bas63 that encodes the two-gene operon, the observed NAD^+^ depletion was less severe, showing that this phage can partially overcome NAD^+^ depletion by SIR2- HerA (Figure 1E).

Considering the pfam domain annotations of the two phage genes, we expected these genes to produce NMN as their end product (Figure 1B). There are two NAD^+^ salvage pathways in *E. coli* that can use NMN to synthesize NAD^+^, either via the enzyme PncC^17^ or via the enzyme NadR^18^. We therefore hypothesized that deletion of *pncC* and *nadR* from the *E. coli* genome would render phage Bas63 sensitive to SIR2-HerA defense despite encoding the two anti-defense genes. However, surprisingly, wild type Bas63 was still able to overcome NAD^+^ depletion-based defense in a Δ*pncC*Δ*nadR E. coli* strain that encoded SIR2-HerA (Figure 1F). These results suggested that the mechanism employed by Bas63 to counter SIR2-HerA defense does not require the NAD^+^ salvage pathway naturally encoded by *E. coli*. These data also implied that the pfam annotations of the two phage genes might be incorrect, and that these proteins synthesize a molecule other than NMN.

### A new two-step biochemical pathway produces NAD^+^ from ADPR and nicotinamide

To gain further insights into the mechanism by which the two phage genes increase NAD^+^ levels during infection, we used untargeted mass spectrometry (MS) to analyze small metabolites in cells infected with the wild type Bas63, and compared these metabolites to those found in cells infected by the mutant phage in which the two genes were deleted. These analyses revealed the presence of two unique molecules with m/z values of 720.010 and 622.034, respectively (in positive ionization mode). These molecules were exclusively observed when the cells were infected with the wild type Bas63 in the presence of the SIR2-HerA defense system, but not in cells infected by the mutant Bas63 phage (Figure 2A, B and Figure S1A, B). The molecules were also not observed in control cells lacking the system that were infected by wild type Bas63, suggesting that these unique molecules are only formed in the presence of the defense system, and only when the phage encodes the two anti-defense genes (Figure 2A, B and Figure S1A, B). Tandem mass spectrometry analysis (MS/MS) of the 720.010 molecule revealed fragments conforming with a pyrophosphorylated form of ADPR (ADPR pyrophosphate, or ADPR-PP), a molecule that, to our knowledge, was not described previously in biological systems (Figures 2C, S1C). MS/MS analysis of the second mass suggested it to be a likely product of ADPR-PP spontaneous hydrolysis, ADPR-cyclic-phosphate (ADPR-cP) (Figure S1D).

**Figure 2.**
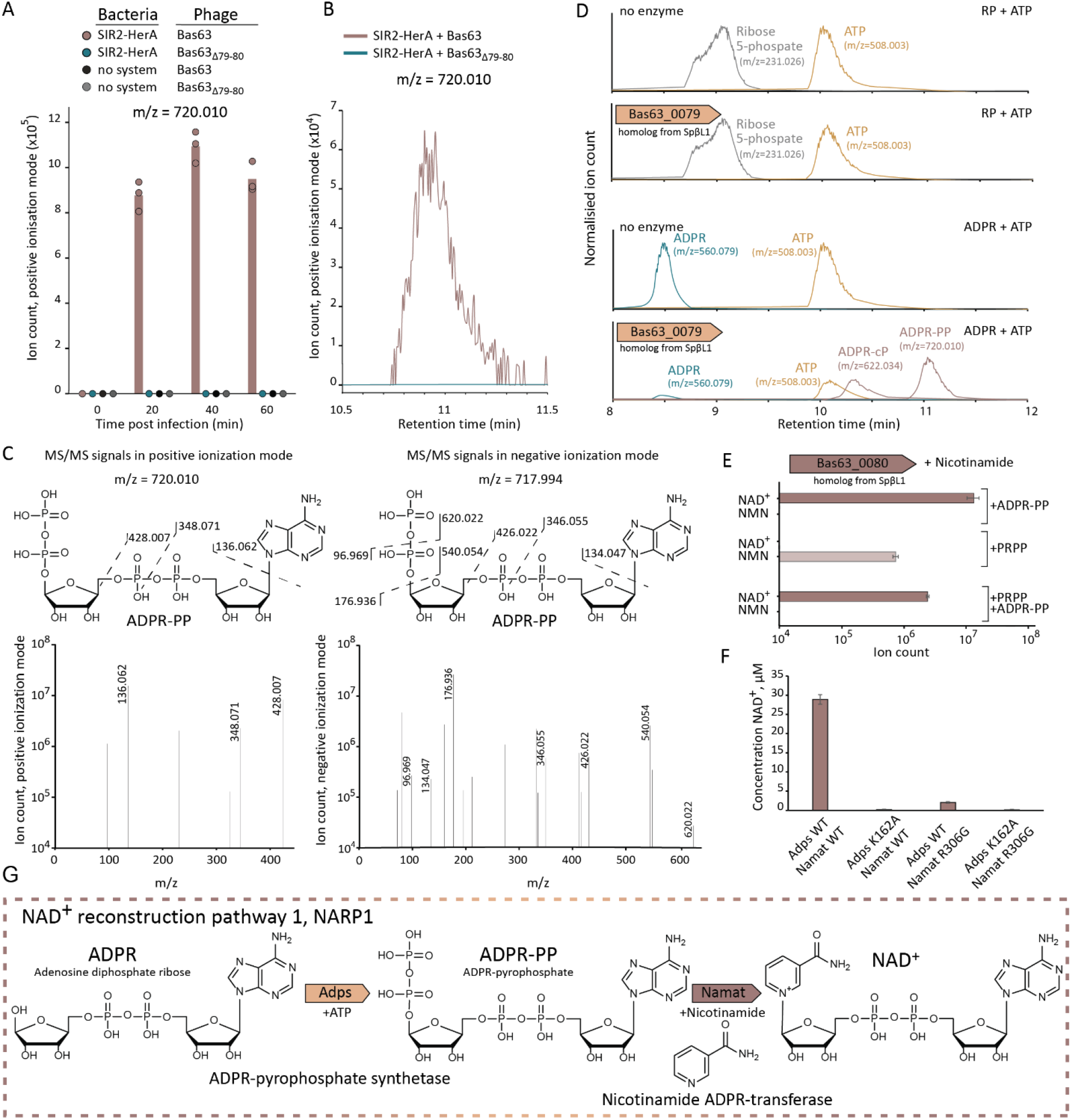
A two-enzyme pathway reconstitutes NAD^+^ from ADPR and nicotinamide. **A**. A unique molecule with an m/z value of 720.010 appears in SIR2-HerA-expressing cells infected by Bas63. Cells were infected at MOI=10. Presented are LC-MS ion counts data, bars represent the mean area under the curve (AUC) of three experiments, with individual data points overlaid. **B**. Extracted mass chromatograms of ions with an m/z value of 720.010 (positive ionization mode) and retention time of 11.0 min, from a lysate of SIR2-HerA cells 20 minutes post infection by phage Bas63 or Bas63_Δ79-80_. Chromatograms of the same molecule in negative ionization mode are presented in Figure S1C. **C**. MS/MS fragmentation spectra of the molecule with the m/z value of 720.010, in positive and negative ionization modes. The hypothesized structure of the molecule and the corresponding MS/MS fragments are presented. **D**. A homolog of Bas63_0079 was purified from phage SpβL1 and incubated either with ribose-5-phosphate or with ADPR in the presence of ATP. Shown are LC-MS analyses of the enzymatic reactions. Peak intensities of each compound were normalized to the signal of the corresponding standard sample (Methods). Representative chromatograms of three replicates are presented. **E**. A homolog of Bas63_0080 was purified from phage SpβL1, and incubated either with ADPR-PP, PRPP or a mixture of ADPR-PP and PRPP, in the presence of nicotinamide. Shown are data for the molecules NAD^+^ and NMN measured via LC-MS. Bars represent the mean area under the curve (AUC) of three experiments, error bars represent standard deviation. **F**. WT Adps and WT Namat, as well as active site mutated versions, were incubated with nicotinamide, ADPR and ATP. NAD^+^ concentration was determined using the NAD/NADH-Glo assay. Bar graphs represent average of three replicates, error bars represent standard deviation. **G**. Schematic of the NARP1 NAD^+^ reconstitution pathway.

The possible appearance of ADPR-PP in SIR2-HerA-expressing cells infected by Bas63 led us to hypothesize that the phage gene annotated as *prs* (the first gene in the two-gene phage operon, Figure 1A) does not encode a phosphoribosylpyrophosphate synthetase enzyme, but rather an enzyme that pyro-phosphorylates ADPR. Under this hypothesis, instead of adding two phosphates to ribose-5-phosphate to generate PRPP, the phage enzyme would catalyze the addition of two phosphates to ribose moiety of ADPR to generate ADPR-PP. PRPP is known to be easily spontaneously hydrolyzed to form 5-phosphorylribose 1,2-cyclic phosphate (PRcP)^19^. Likely, the ADPR-PP analog could also undergo similar spontaneous hydrolysis, leading to the formation of ADPR- cyclic-phosphate and possibly explaining the appearance of the ion with the 622.034 m/z value (Figure S1A and B).

To further examine the enzymatic activity of the putative ADPR-PP-generating phage enzyme, we attempted to express and purify it for *in vitro* analyses. As the protein from Bas63 did not purify well, we instead used a homologous operon from phage SpβL1^20^ from which we were able to obtain a purified protein. Incubation of the purified protein with PRPP and ATP did not yield any measurable product, further supporting that this enzyme is not Prs (Figure 2D). However, incubation of the protein with ADPR and ATP resulted in the formation of the same two molecules detected in our analysis of cellular metabolites during infection *in vivo* (Figure 2D).

To establish the identity of the enzyme products, we treated the products of the enzymatic reaction with NudC, an enzyme of the NUDIX family that cleaves internal nucleoside- linked phosphate-phosphate bonds. As expected from the structure of ADPR-PP, incubation of the molecule suspected as ADPR-PP with NudC resulted in the formation of PRPP, and similarly, NudC treatment of the molecule suspected as ADPR-cP yielded PRcP as a product (Figure S2A). Treatment with apyrase, an enzyme which degrades terminal diphosphates into monophosphates, resulted in the formation of ADPR monophosphate (ADPR-P) from ADPR-PP (Figure S2B). Combined together, these results strongly suggest that the purified phage enzyme catalyzes the addition of pyrophosphate to ADPR in an ATP-dependent manner. To our knowledge, pyrophosphorylation of ADPR is an enzymatic reaction never previously described. We named the phage enzyme **AD**PR-**P**P **s**ynthetase (Adps).

Based on its sequence annotation, the second protein in the phage operon was expected to transfer a nicotinamide molecule to PRPP, replacing the high-energy pyrophosphate moiety with nicotinamide to generate NMN (Figure 1B). Given that the first enzymatic reaction in this pathway generates ADPR-PP rather than PRPP, we hypothesized that the second enzyme would use ADPR-PP as a substrate, and conjugate the nicotinamide instead of the pyrophosphate to directly generate NAD^+^. To test this hypothesis, we expressed and purified this second enzyme from the operon of phage SpβL1, and incubated the purified protein with ADPR-PP and nicotinamide. This resulted in the formation of NAD^+^, confirming our hypothesis (Figure 2E). To our knowledge, this reaction, too, was not previously described in other biological systems. We named the second phage enzyme **N**icotin**am**ide **A**DPR-**t**ransferase (Namat).

When incubated with PRPP and nicotinamide, the phage Namat enzyme was capable of producing NMN; but when exposed to a mixture of PRPP and ADPR-PP together with nicotinamide, only NAD^+^ was detected as a product, suggesting a preference for this enzyme to use ADPR-PP as a substrate (Figure 2E). Incubating a mixture of Adps and Namat with ADPR, nicotinamide and ATP resulted in the production of NAD^+^ *in vitro* (Figure 2F). The two-enzyme mixture was not able to efficiently produce NAD^+^ if the predicted active site of Adps was impaired by substituting K162 to alanine, or if the predicted active site of Namat was modified by a R306G substitution, further confirming that both enzymatic activities are essential for NAD^+^ reconstitution (Figure 2F). Together, our results reveal a phage-encoded novel two-step enzymatic pathway that can reconstitute NAD^+^ from ADPR and nicotinamide in an ATP-dependent manner (Figure 2G). We name this pathway NAD^+^ reconstitution pathway 1 (NARP1).

### Phage-mediated NAD^+^ reconstitution overcomes multiple defense systems

NAD^+^-depleting defense systems that rely on SIR2, TIR or SEFIR domains all break NAD^+^ into ADPR and nicotinamide to achieve NAD^+^ depletion^6-8^. Therefore, a pathway that can use these same products to re-build NAD^+^ would be an efficient solution for the phage to counter NAD^+^ depletion by any of these defense systems. To test this hypothesis, we examined the effect of the NARP1 pathway on four additional defense systems: Type I Thoeris, DSR1, DSR2, and SEFIR. Type I Thoeris is a two-gene system that contains an effector with a SIR2 domain, which was shown to deplete NAD^+^ in response to phage infection^7^. DSR1 and DSR2 are large defensive proteins, each of which activates an N-terminal SIR2 domain to deplete NAD^+^ when a phage is detected by the C-terminal domain of the protein^6^; and SEFIR is a single-protein defense system that depletes NAD^+^ via its N-terminal SEFIR domain^8^. Co-expression of the NARP1 pathway in *Bacillus subtilis* with any of these systems abolished the system’s ability to defend against *Bacillus* phages, confirming the generality of NARP1 in countering multiple defense systems (Figure 3A). Point mutation in the active sites of either of the two proteins in the NARP1 pathway impaired its ability to efficiently counter defense (Figure 3A).

**Figure 3.**
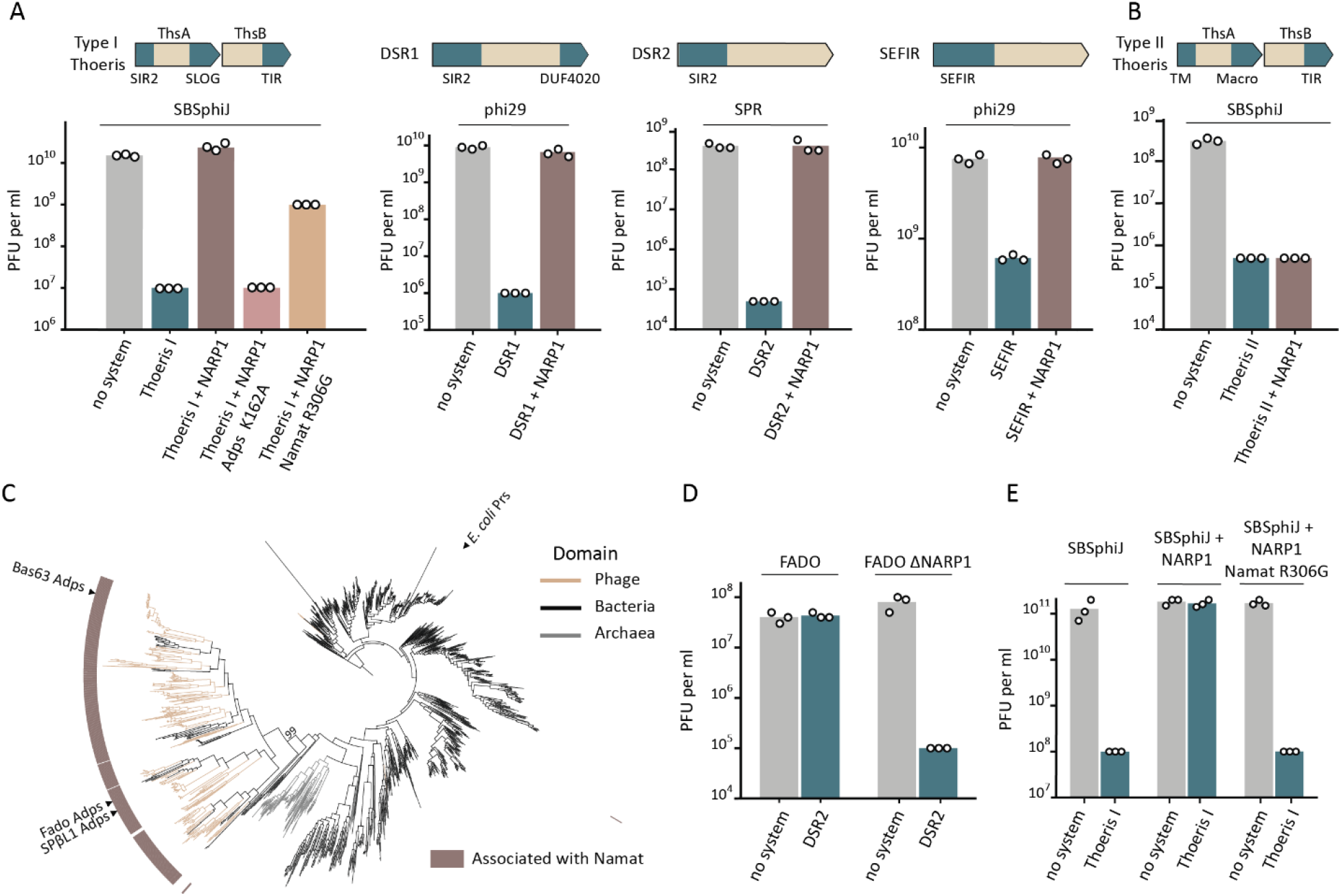
NARP1 is a phage-specific pathway that overcomes multiple NAD^+^ depletion defense systems. **A**. Anti- defense effect of NARP1 co-expressed in *B. subtilis* together with diverse NAD^+^-depleting defense systems. Data represent PFU per ml of *Bacillus* phages infecting control cells (no system), cells expressing the respective defense systems and cells co-expressing the defense system and NARP1. For Type I Thoeris, results for co-expression of mutated versions of NARP1 are also presented. **B**. NARP1 does not overcome the anti-phage defense conferred by type II Thoeris, a defense system that does not deplete NAD^+^ as part of its mechanism. Data presented are as in panel A. **C**. Phylogenetic analysis of the Adps/Prs protein family in phages, bacteria and archaea. Genes were marked as associated with Namat if a Namat homolog was detected in the genomic vicinity (up to 10 genes away from the Prs/Adps-encoding gene). Prokaryotic and viral protein sequences were clustered separately based on identical length and >90% identity and a representative sequence of each cluster was used to build the tree (see Methods). Ultrafast bootstrap value^23^ is shown for the Adps clade. **D**. Deletion of the NARP1 operon sensitizes the FADO phage to the DSR2 defense system. Data represent PFU per ml of FADO and FADO_ΔNARP1_ phages infecting control cells (no system) and cells expressing the DSR2 defense system. **E**. Knock-in of NARP1 into phage SBSphiJ results in a phage that overcomes the type I Thoeris defense system. Data represent PFU per ml of WT SBSphiJ, SBSphiJ with NARP1 knock-in, and SBSphiJ knocked in for NARP1 with an active site mutation in Namat. Phages were used to infect control cells (no system) and cells expressing the type I Thoeris defense system. In panels A, B, D and E, bar graphs represent average of three independent replicates, with individual data points overlaid. In each panel, the phages that were used are indicated above the respective bar graphs.

As a control for these experiments, we used type II Thoeris, a defense system that functions similar to type I Thoeris, but in which the SIR2 domain is replaced by two transmembrane helices that are thought to induce membrane permeability when the system detects phage infection, causing premature cell death independent of NAD^+ 21^. Expression of NARP1 with type II Thoeris did not abolish defense, further supporting that NARP1 can counter only bacterial defenses that rely on NAD^+^ depletion (Figure 3B).

### The NAD^+^ reconstitution pathway is phage-specific

Given the homology between the phage enzyme Adps and the bacterial Prs enzyme, we sought to study the evolutionary relationship between the two enzymes and their taxonomic distribution across the phylogenetic tree. For this, we searched a database of ∼4,000 finished bacterial and archaeal genomes^8^ and a database of ∼20,000 complete phage genomes^22^ for homologs of Prs and Adps. A phylogenetic analysis of the detected homologs revealed that Adps proteins localize to a well-supported clade that is separated from the majority of prokaryotic Prs homologs, and that appears almost exclusively in phage genomes (Figure 3C). In a minority of cases, sequences in the Adps clade were present in bacterial genomes, and in some, but not all of these cases, Adps was within prophages or other mobile genetic elements integrated in the bacterial genome (see Discussion; Table S1).

We next examined the genomic neighborhood of genes encoding Prs and Adps homologs. For the vast majority of proteins in the Adps clade (97%) we detected a homolog of Namat in the immediate vicinity of the *adps* gene, confirming the functional association between Adps and Namat within the NARP1 pathway (Figure 3C). In contrast, bacterial *prs* genes were only very rarely present next to a gene encoding a Namat homolog. Altogether, our analyses suggest that Adps proteins form a subfamily within the phosphoribosylpyrophosphate synthetase family of proteins, and that such Adps proteins evolved to utilize ADPR as a substrate instead of ribose-5-phosphate as part of the NARP1 pathway.

We detected the NARP1 pathway encoding Adps and Namat in 859 sequenced phage genomes, representing 4.3% of the phages in the set we analyzed (Table S1). To assess the generality of our findings we asked whether additional homologs of this NAD^+^ reconstitution pathway also endow phages with the ability to evade NAD^+^ depletion- mediated defense. For this, we examined NARP1 from phage Fado, a lytic phage that we previously isolated^8^, and generated a mutant of Fado where the two genes encoding NARP1 were deleted. The mutant phage could not replicate in cells expressing the defense system DSR2, while the wild type Fado phage overcame defense (Figure 3D). To further test whether NARP1 can protect phages from the effects of NAD^+^ depletion independently of other phage factors, we engineered NARP1 from SpβL1 into phage SBSphiJ, a phage that is naturally sensitive to the type I Thoeris defense system. We found that the engineered phage can now overcome the type I Thoeris system (Figure 3E), whereas a phage engineered with mutated NARP1 could not overcome Thoeris defense. Our data confirm the broad function of the viral NARP1 pathway in counteracting NAD^+^ depletion-based bacterial defenses.

### A second pathway in phages overcomes NAD^+^-depleting defense systems

We next searched for homologs of the NARP1 Namat proteins in the set of ∼20,000 phage genomes. While the majority (∼79%) of Namat proteins were found in phages that also encode Adps, we observed that ∼20% of homologs were encoded in phages that lack any homolog of Adps. A phylogenetic analysis showed that these homologs organize into a clade distinct from the clade encoding the NARP1-associated Namat proteins, suggesting that this clade encodes proteins that have sequence homology to NARP1 Namat enzymes but may have a different enzymatic function (Figure 4A; Table S2).

**Figure 4.**
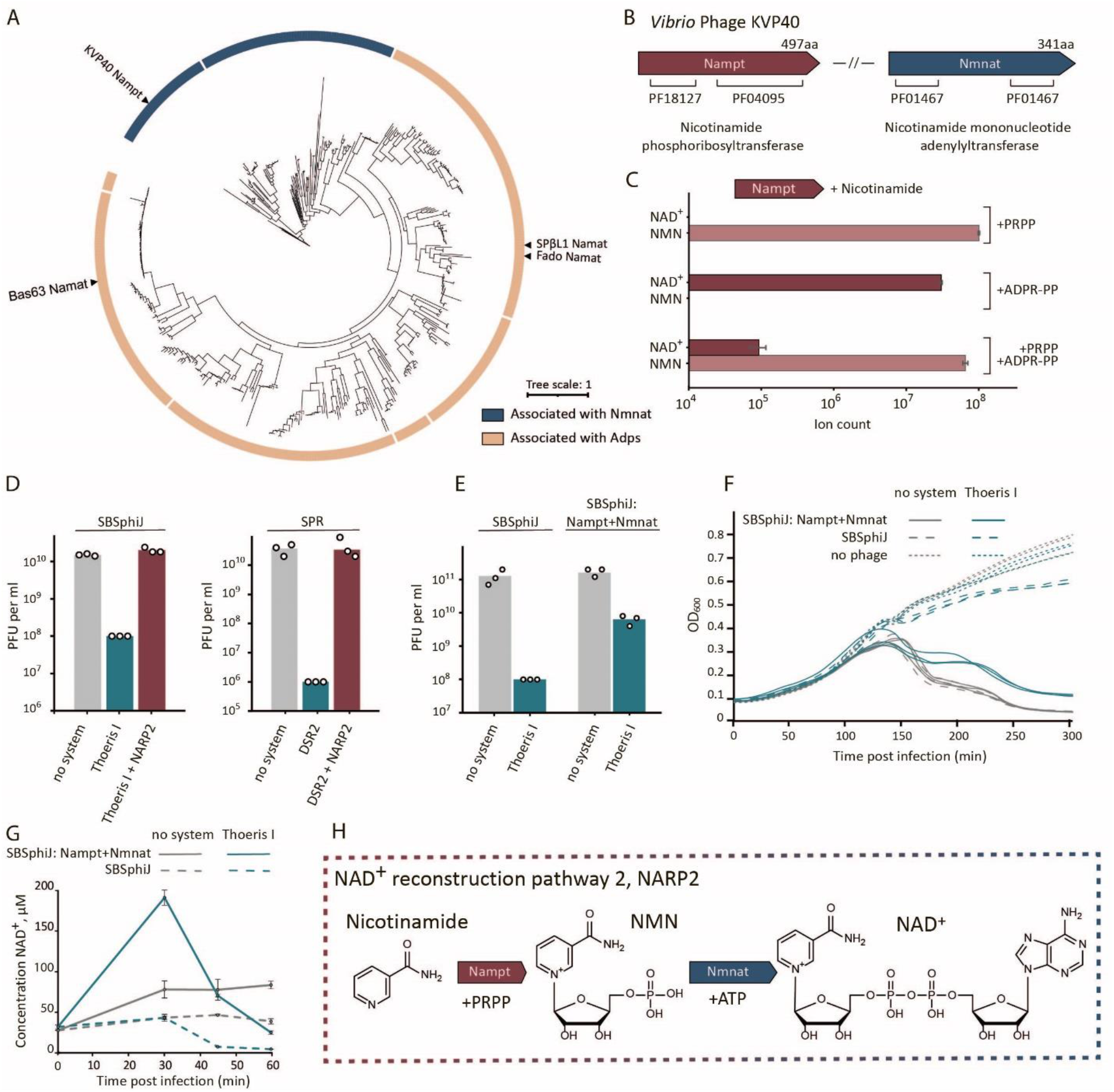
NARP2 is a second NAD^+^ reconstitution pathway allowing phages to overcome bacterial defense. **A**. Phylogenetic analysis of the Namat/Nampt protein family in phage genomes. Outer ring indicates whether the phage genome encoding the respective gene also encodes nicotinamide mononucleotide adenylyltransferase (Nmnat, blue) or whether the gene is associated with Adps (beige color) as defined in Figure 3C. Protein sequences were clustered based on identical length and >90% identity and a representative sequence of each cluster was used to build the tree (see Methods). **B**. Domain annotations of the NAD^+^ salvage proteins from *Vibrio* phage KVP40. **C**. Nampt from *Vibrio* phage KVP40 was purified and incubated *in vitro* with either PRPP, ADPR-PP, or a mixture of ADPR-PP and PRPP, in all cases in the presence of nicotinamide. The products NAD^+^ and NMN were measured by LC- MS. Data represent the mean area under the curve (AUC) of three experiments, error bars represent standard deviation. **D**. NARP2, when co-expressed with NAD^+^-depleting defense systems, inhibits anti-phage defense. Data represent PFU per ml of phages infecting control cells (no system), cells expressing the indicated defense systems and cells co-expressing the defense system and NARP2. **E**. Knock-in of NARP2 into phage SBSphiJ results in a phage that partially overcomes type I Thoeris defense. Data represent PFU per ml of phages SBSphiJ and SBSphiJ+NARP2 infecting control cells (no system) and cells expressing type I Thoeris. For panels D and E, bar graphs represent average of three independent replicates with individual data points overlaid. Phages used for the infection assays are indicated above the graph. **F**. Liquid culture growth of *B. subtilis* cells expressing the type I Thoeris defense system, or control cells without defense genes (no system). Cells were infected by phages SBSphiJ and SBSphiJ with the NARP2 pathway knocked in. Infection was performed at MOI=0.1. Data from three replicates are presented as individual curves. **G**. NAD^+^ concentration in lysates extracted from cells expressing type I Thoeris or control strain (no system). Cells were infected either with phage SBSphiJ or with SBSphiJ knocked in with the NARP2 pathway, at MOI=10. Cells were collected before infection (0 min) and 30, 45 or 60 minutes after infection. NAD^+^ concentration was determined using the NAD/NADH-Glo assay. The experiment was performed in three replicates, error bars represent standard deviation. **H**. Schematic of the enzymatic reactions of NARP2^24^.

One of the proteins in the new clade is encoded in *Vibrio* phage KVP40 (Figure 4A, 4B). This protein was previously shown to have a nicotinamide phosphoribosyltransferase activity (Nampt), capable of producing NMN from PRPP and nicotinamide^24^. To test whether this protein is also capable of producing NAD^+^ from ADPR-PP, we purified the protein and incubated it with either PRPP, ADPR-PP, or both PRPP and ADPR-PP, all in the presence of nicotinamide and ATP. Our data reproduced the previous observation that the protein has nicotinamide phosphoribosyltransferase enzymatic activity^24^. Although we could observe the production of NAD^+^ from ADPR-PP, a reaction in which both ADPR-PP and PRPP were supplied as substrates to the purified protein showed that this enzyme prefers PRPP as a substrate (Figure 4C).

It was previously shown that a second gene in *Vibrio* phage KVP40 encodes a nicotinamide mononucleotide adenylyltransferase (Nmnat), capable of generating NAD^+^ by conjugating AMP to NMN^24^ (Figure 4A). Moreover, it was shown that the *Vibrio* phage KVP40 Nmnat, together with the nicotinamide phosphoribosyltransferase from the same phage, can generate NAD^+^ from ribose-5-phosphate, nicotinamide and ATP^24^. Given that these two proteins represent an NAD^+^ reconstitution pathway that functions via enzymatic reactions different than those encoded by the NARP1 pathway, we hypothesized that the KVP40 pathway, too, can save phages from the effects of NAD^+^-depleting defense systems.

To test this hypothesis we co-expressed these two genes in cells expressing either a type I Thoeris or DSR2, and infected these cells by phages known to be blocked by the respective defense systems. Our data show that the two genes from KVP40 completely abolished the activity of both Thoeris and DSR2 (Figure 4D). We next integrated these two genes from phage KVP40 into the genome of phage SBSphiJ, which is naturally blocked by type I Thoeris, and found that the engineered phage was able to overcome Thoeris defense (Figure 4E, 4F).

To further examine whether the two genes supply the phage with NAD^+^ despite the activity of the defense system, we measured NAD^+^ levels during infection, and detected a significant increase in NAD^+^ levels in cells infected by the SBSphiJ phage engineered to express the two genes from KVP40 (Figure 4G). The increase of NAD^+^ levels was also observed in SBSphiJ-infected *B. subtilis* cells that did not encode the defense system, suggesting that this pathway generates NAD^+^ regardless of the activity of NAD^+^-depleting defense systems (Figure 4G), corroborating previous reports on the *Vibrio* KVP40 phage that naturally encodes these genes^24^.

Our data show that almost all phages (94%) in which we detected a homolog of the KVP40 Nampt enzyme also encode a homolog of Nmnat, providing further support that these two proteins are functionally linked and form an NAD^+^-reconstitution pathway (Figure 4H; Table S2). Our data suggest that the function of this pathway is to counter the activity of NAD^+^-depleting defense systems. We name this pathway NAD^+^ reconstitution pathway 2 (NARP2), and find that NARP2 is encoded in ∼1.1% of sequenced phage genomes (Table S2). We were not able to find any phage that encodes both NARP1 and NARP2 (Table S2).

## Discussion

Phages are known to encode multiple proteins to counter bacterial defenses^20,25,26^. In most cases, each anti-defense phage protein inhibits only a narrow range of defense systems. Most anti-CRISPR proteins, for example, bind and inhibit only a specific subtype of CRISPR-Cas^27^, and Apyc1, a phage protein that degrades immune signaling molecules, only inhibits the Pycsar system^28^. The phage anti-defense strategy we describe in the current study is unique because it counters the consequences of the immune action rather than components of the defense system itself. Thus, it allows phages to evade a wide variety of systems that deplete NAD^+^, including SIR2-HerA^6^, DSR1^6^, DSR2^6^, and type I Thoeris^7^, and is likely to also overcome variants of CBASS^11^, Pycsar^12^, AVAST^10^, and pAgo^6,9^ that have effectors that deplete NAD^+^.

While both NARP1 and NARP2 rebuild NAD^+^ as their final products, they differ in their substrates. NARP1 uses ADPR and nicotinamide, the direct products of NAD^+^ cleavage by immune effector domains such as TIR, SIR2 and SEFIR^6-8^, and necessitates only ATP in addition to the cleavage products for the production of NAD^+^. This pathway will therefore not come into action unless the phage infects a cell that contains an NAD^+^- depleting system, and unless the system actively depletes NAD^+^ in attempt to protect against the infecting phage. As ADPR is normally not present in substantial quantities in uninfected *B. subtilis* cells^6^, this pathway likely inflicts minimal metabolic costs to the phage when infecting cells that do not encode an NAD^+^-depleting defense system. Indeed, we did not observe production of excessive NAD^+^ when NARP1-containing phages infected cells that lack a defense system (Figure 1E). In contrast, phages encoding NARP2 caused cells to produce excessive NAD^+^ during infection even when the cells did not contain an NAD^+^-depleting defense system, because NARP2 uses PRPP as its starting substrate (Figure 4G). Thus, NARP2 may inflict a more severe metabolic cost to the phage, especially as it consumes PRPP, which is essential for nucleotide synthesis^29^. The possibly higher metabolic cost inflicted by NARP2 may explain why it is rarer in phage genomes as compared to NARP1.

Since at least 7% of all bacteria whose genomes were sequenced encode an NAD^+^- depleting defense system^8^, it is not surprising that NARP1 and NARP2 together are present in 5.4% of sequenced phages. A recent analysis of phage genomes shows that in addition to NARP1 and NARP2, phages can encode other genes whose annotations suggest involvement in NAD^+^ salvage^15^. For example, some phages encode homologs of NadR, an enzyme known to produce NAD^+^ from nicotinamide riboside and ATP^15^. Phage NadR -encoding genes are usually present in the same operon as a gene annotated as encoding PnuC, a transporter specific for nicotinamide riboside, and it is conceivable that this operon would form yet another pathway allowing phages to produce NAD^+^ via available metabolites (Figure S3). Thus, the actual fraction of phages encoding an NAD^+^ reconstitution pathway may be even larger than the >5% we recorded.

While NARP1 is preferentially encoded in phages, some bacterial genomes seem to encode it not in the context of a prophage or a mobile genetic element (Table S1). ADPR was shown, in a minority of bacteria, to be present at detectable concentrations^30^ and it was demonstrated that this metabolite may be involved in regulation of transcription in some bacteria^31^. It is therefore possible that some bacteria have adopted NARP1 for housekeeping or regulatory functions not related to the phage-bacteria conflict; future studies will be necessary to determine the role of NARP1 in this context.

In recent years, multiple bacterial immune systems were shown to degrade essential metabolites as a measure of anti-phage defense, representing a general strategy of depriving the phage from an essential metabolite and limiting its propagation. In addition to NAD^+^ depletion, metabolite-depleting defense systems include dCTP deaminases^32^ and dGTPases^32^ that degrade deoxynucleotides, and ATP nucleosidases that deplete ATP in infected cells^13^. It is possible that, similar to NAD^+^ reconstitution pathways, phages may also encode pathways that rebuild deoxynucleotides or ATP from the degradation products generated by the respective defense system. As depletion of deoxynucleotides and ATP was also shown to be utilized by animal cells as an antiviral measure^13,33^, future studies may reveal that viruses infecting animals may also use metabolite reconstitution as a counter-defense strategy.

## Materials and Methods

### Strains and growth conditions

*E. coli* K-12 BW25113, DH5a and BL21 (DE3) were grown in MMB media (lysogeny broth (LB) supplemented with 0.1 mM MnCl_2_ and 5 mM MgCl_2_) at 37°C with 200 rpm shaking or on solid 1.5% LB agar plates. Ampicillin 100 μg/ml, chloramphenicol 30 μg/ml or kanamycin 50 μg/ml were added when necessary for plasmid maintenance. *E. coli* DH5a (NEB) was used for cloning, BL21 (DE3) for protein purification and BW25113 for experiments with phages. *B. subtilis* strain BEST7003 (obtained from Mitsuhiro Itaya of Keio University, Japan) was grown in MMB at 25°C or 30°C, spectinomycin 100 μg/mL and/or chloramphenicol 5 μg/ml were added when needed. All chemicals were obtained from Sigma Aldrich unless stated otherwise. All phages used in the study were amplified from a single plaque at 37°C (except for FADO, which was amplified at 25°C) in *B. subtilis* BEST7003 culture in MMB until the culture collapsed. A list of all plasmids, strains and phages used in this study can be found in Supplementary Table S3.

### Plasmid construction and transformation

DNA amplification for cloning was made by KAPA HiFi HotStart ReadyMix (Roche) according to manufacturer instruction. All primers were obtained from Sigma Aldrich. Supplementary Table S4 lists all primers used in this study.

For site-directed mutagenesis, the whole plasmid was amplified by back-to-back primers containing mutations at the primer 5’-end. PCR product was then directly used for circularization with KLD enzyme mix (NEB) according to manufacturer instructions, and used for transformation. For cloning of large fragments, PCR products with 20-nucleotides overlaps were generated and treated with FastDigest DpnI (ThermoFisher) restriction enzyme for 30 min at 37°C. Fragments were then Gibson assembled by NEBuilder HiFi DNA Assembly Master Mix (NEB) according to manufacturer’s instructions and used for transformation into DH5a (NEB #C2987H). Single colonies were checked by PCR and plasmids were validated by a plasmid sequencing service (Plasmidsaurus).

Verified plasmids were transformed into *E. coli* using the standard TSS protocol^34^ or to *B. subtilis* using MC media as previously described^35^. The KVP40-NARP2 operon (Table S3) could not be transformed into *E. coli* because of toxicity, and hence the Gibson assembly product was directly transformed into *B. subtilis* using the MC media protocol. Integrations of the NARP1 and NARP2 constructs into *B. subtilis* strains were confirmed by whole-genome sequencing.

To design active site mutations of Adps and Namat, we aligned their sequences with previously-studied homologs. SpβL1 Adps is similar to *B. subtilis* Prs (27% sequence identity), in which Lys197 was previously demonstrated biochemically and structurally to be essential for the enzymatic activity^36,37^. Therefore, Lys162 of Adps from SpβL1, corresponding to Lys197 of *B. subtilis* Prs was mutated to alanine by site-directed mutagenesis. SpβL1 Namat shares 33% sequence identity with the *Homo sapiens* Nampt protein, the active site of which was studied biochemically and structurally^36^. A Arg306 to Gly mutation was designed in the SpβL1 Namat based on the homologous residue Arg311 shown to be important for the activity of the human Nampt^36^. Primers for site-directed mutagenesis of the both genes are in the Supplementary Table 4.

### Deletion of NARP1 operons from Bas63 and FADO phages

The NARP1 operon from Bas63 was deleted using Cas13a as previously described^38^. A gRNA complementary to the beginning of the coding region of gene 79 was cloned to the plasmid pBA559 treated with BsaI. *E. coli* DH5a was transformed with this plasmid and then infected by Bas63 in two consecutive rounds with MOI∼1 in 5 ml of MMB media. Then, 100 µl of lysate was spread on a plate with the same strain (DH5a without plasmid was used as a control). Several plaques were collected and screened by PCR. Deletion of the genomic region starting from codon 28 of gene 79 and ending in coding 302 of gene 80 was confirmed by whole genome sequencing of the phage genome.

To delete the NARP1 region from FADO phage, lysogenic bacteria were first prepared by infecting *B. subtilis* strain BEST7003 cells with the FADO phage (MOI 0.1). When cells started to grow again after lysis, the culture was spread onto an MMB-agar plate. Individual clones were selected and checked for lysogeny by incubation with Mitomycin C (0.5 μg/ml) followed by phage titer counting in the supernatant. Lysogenic bacteria were transformed with a non-replicating in *B. subtilis* plasmid (pJmp3) encoding a spectinomycin resistance gene flanked by 1 kb regions identical to the DNA flanking NARP1 in the FADO genome. Following homologous recombination, spectinomycin-resistant lysogens were selected on a plate with spectinomycin and checked by PCR. The modified FADO prophage was induced using Mitomycin C (0.5 μg/ml). Whole-genome sequencing was performed to verify the sequence of the modified phage.

### Knock-in of intact and mutated NARP1 and NARP2 operons into SBSphiJ

Insertion of DNA into the SBSphiJ genome by means of homology recombination was demonstrated previously to be achievable via homologous recombination^39^. Here we used the same region in the SBSphiJ genome previously used to knock-in the gene *tad1*^39^, and the same flanking regions, to integrate the NARP1 and NARP2 operons into the SBSphiJ genome. Plasmids with intact and mutated NARP1 and Gibson assembly product for NARP2, flanked by 1 kb regions identical to the DNA flanking Tad1 in the SBSphiJ7 genome, were transformed to *B. subtilis* BEST7003 and these strains were infected by SBSphiJ with MOI 0.1 for generating recombined phages. A gRNA complementary to unmodified SBSphiJ was cloned into the previously published plasmid pGad2-Cas13a^20^ to generate pCas13a-gRNA-SBSphiJ. *B. subtilis* BEST7003, transformed with this plasmid was used for selection against unmodified SBSphiJ phages in order to retain only phages where homology recombination took place. The selection was made on an agar plate in the presense of 0.2% xylose and several plaques were tested by PCR for the presence of intact and mutated NARP1 and NARP2. Whole-genome sequencing was performed to verify the sequence of the modified phage.

### Generation of Δ*pncC*Δ*nadR E*. *coli* strain by P1 transduction

BW25113 Δ*pncC* strain from the Keio collection^40^ was transformed with the pCP20 plasmid to eliminate the resistance cassette^40^. Kanamycin sensitive clones were selected and checked by PCR. BW25113 Δ*nadR* from the Keio collection was infected with P1 and used as a donor for transduction, which was made according to the standard protocol^41^. Double-knockouts were selected on kanamycin and checked by PCR, positive clones were grown twice on LB media with 5 mM sodium citrate to eliminate the residual phage. Sequence of the Δ*pncC*Δ*nadR E. coli* was validated by a genome sequencing service (Plasmidsaurus).

### Plaque assays

Phage titer was determined as described previously^42^. 300 µl of the overnight bacterial cultures were mixed with 30 ml of melted MMB 0.5% agar, poured on 10 cm square plates and left to dry for 1 h at room temperature. IPTG was added to a concentration of 1mM to induce NARP1 and NARP2 from the *B. subtilis* genome. Tenfold dilutions of phages were prepared in MMB and 10 µl of each dilution was dropped onto the plates. Plates were incubated overnight at 25°C (DSR1, DSR2, SEFIR, type II Thoeris), 30°C (type I Thoeris) and 37°C (SIR2-HerA). Plaque-forming units were counted the next day.

### Liquid infection assays

Overnight bacterial cultures were diluted in MMB and grown until reaching an optical density at 600nm (OD_600_) of 0.3. Then, 180 µl of cultures were transferred to a 96-well plate and infected with 20 µl of phages at various MOIs. Culture growth was followed by OD_600_ measurements every 10 min on a Tecan Infinite 200 plate reader. Cells expressing the SIR2-HerA system were grown at 37°C, while those expressing the type I Thoeris system were grown at 30°C.

### Proteins purification

Plasmids were transformed into BL21 (DE3) cells, and cells were grown in MMB media at 37°C until mid-log phase (OD_600_∼0.5), then IPTG was added to a concentration of 0.5 mM and cells were further grown for 3 additional hours, centrifuged and flash-frozen. His-tagged proteins were purified by NEBExpress® Ni-NTA Magnetic Beads (NEB) according to the manufacturer’s protocol. Proteins were transferred to 20 mM Tris (pH 8), 200 mM NaCl, 5 mM DTT and 5%(v/v) glycerol by five cycles of filtration through 10 kDa Amicon filters (Merck Millipore), and finally concentrated to 0.5 mg/ml.

### Enzymatic activity of purified proteins

Reactions were performed in 50 mM Tris (pH 7.5), 12 mM MgCl_2_, BSA 0.02% (w/v), D-Ribose 5-phosphate (Sigma 83875), 5-Phospho-D-ribose 1-diphosphate (Sigma P8296), adenosine 5’-diphosphoribose (Sigma A0752) and nicotinamide were added to 1 mM when indicated. ATP was added in excess to a concentration of 5 mM in all reactions, except for the one presented in Figure 2D, where ATP concentrations were 1 mM (equimolar). Adps was added at 0.5 µM, Namat and Nampt in all reactions were at 0.2 µM. Reactions were incubated for 60 min at 30°C and then stopped by adding methanol 50% and diluting 10 times 50% methanol/50% 0.1 M Na-phosphate buffer, pH 8.0. Products and substrate molecules were analyzed by LC-MS or with NAD/NADH-Glo kit (Promega).

### Cell lysates preparation

*E. coli* BW25113 cells carrying a plasmid with the SIR2-HerA system or an empty plasmid were grown at 37 °C, 200 rpm until reaching an OD_600_ of 0.3 in 250 ml of MMB. Cells were then infected with Bas63 and Bas63_Δ79-80_ with MOI=10. Samples were collected before infection and 20, 40 and 60 minutes after infection. At each time point, 50 ml of cells were centrifuged for 10 minutes at 25°C, 4000 *g* for 10 min to pellet the cells and 600 µl of 50% methanol/50% 0.1 M Na-phosphate buffer, pH 8.0 were immediately added to stop further NAD^+^ degradation. Resuspended cells were frozen in liquid nitrogen and stored at -80°C. To extract metabolites, resuspended cells were transferred to FastPrep Lysing Matrix B in a 2 ml tube (MP Biomedicals, no. 116911100) and lysis was achieved as described previously^39^. Lysates were analyzed by LC-MS or with the NAD/NADH-Glo kit (Promega). *B. subtilis* strain BEST7003 expressing type I Thoeris or control strain without defense system were grown at 30 °C, 200 rpm until reaching an OD_600_ of 0.3 in 250 ml of MMB and then infected with SBSphiJ and SBSphiJ+NARP2 with MOI=10. Samples were collected before infection and 30, 45 and 60 minutes after infection. Next processing was done exactly like for *E. coli* samples described above.

### Enzymatic assay for NAD^+^ measurements

The NAD/NADH-Glo (Promega) kit was used for NAD^+^ level measurement. The lysate or *in vitro* reactions were diluted 1:100 in 0.1 M Na-phosphate buffer, pH 8.0. Reactions were performed in a volume of 10 µl (5 µl of sample + 5 µl of reaction mix) according to manufacturer’s protocol and luciferase signal was detected using the Tecan Infinite 200 PRO plate reader every 2 minutes for 30 cycles. NAD^+^ concentrations were calculated from the calibration curve using a set of standard NAD^+^ samples with known concentrations.

### Enzymatic treatment of ADPR-PP and ADPR-cP with Apyrase and NudC

For Apyrase treatment, 20 µl of reaction mixture of ADPR, ATP and Adps (Figure 2D) was mixed with 15 µl of water, 4 µl of 10x Apyrase buffer (NEB) and 1 µl of Apyrase (M0398S, NEB), then incubated for 30 min at 30°C. Products were analyzed by LC-MS.

For NudC treatment, 20 µl of reaction mixture of ADPR, ATP and Adps (Figure 2D) was mixed with 13 µl of water, 4 µl of 10x r3.1 buffer (NEB), 2 µl of 100µM DTT and 1 µl of NudC (M0607S, NEB), then incubated for 30 min at 37°C. Products were analyzed by LC-MS.

### Metabolites analysis via LC-MS

Metabolic profiling of the polar metabolites was performed as described previosly^43^ with minor modifications as described below. Briefly, analysis was performed using Acquity I class UPLC System combined with mass spectrometer Q Exactive Plus Orbitrap™ (Thermo Fisher Scientific), which was operated in both positive and negative ionization modes. The LC separation was done using the SeQuant Zic-pHilic (150 mm × 2.1 mm) with the SeQuant guard column (20 mm × 2.1 mm) (Merck). The Mobile phase B was acetonitrile and Mobile phase A was 20 mM ammonium carbonate with 0.1% ammonia hydroxide in DDW: acetonitrile (80:20, v/v). The flow rate was kept at 200 μl/min, and the gradient was as follows: 0-2 min 75% of B, 14 min 25% of B, 18 min 25% of B, 19 min 75% of B, for 4 min, 23 min 75% of B.

For the detection of NAD^+^ and NMN by LC-MS, analysis was performed by two separate injections in positive and negative ionization modes from m/z 75 to 1000 at a mass resolution of 70,000. Peak areas were extracted using MZmine 2 with an accepted deviation of 5 ppm. In figure 2D signals of all substrates were normalized to the signals of the standard samples with the same concentration as used in the reaction. Signals for ADPR-PP and ADPR-cP were calculated from the decrease of used ADPR, because reactions were carried out with an excess ATP. All molecules used as standards were obtained from Sigma: ADPR (A0752), PRPP (P8296), D-Ribose 5-phosphate (83875), ATP (A1852), Nicotinamide (72340) NMN (N3501) and NAD^+^ (N8285).

For detection of ADPR-PP and ADPR-cP, MS/MS spectra collection was performed using the same instrument at a resolution of 17,500. Fragmentation was done through a higher-energy collisional dissociation cell using a normalized collision energy 30. Fragments were extracted using MZmine 2.

### Phylogenetic analyses of Adps and Namat

Homologs of Prs/Adps, Namat and Nmnat were searched in 3,895 finished prokaryotic genomes (including 3,781 bacterial and 114 archaeal genomes)^8^ downloaded in October 2017, as well as in 20,185 phage genomes from the INPHARED database downloaded on May 1st 2023^22^. For the Prs/Adps analysis, the *E. coli* Prs protein, phage Bas63 Adps and phage SpβL1 Adps were used as queries the Namat analysis, the Namat protein sequences from phages Bas63, SpβL1 and the Nampt from KVP40 were used. Nmnat was searched using the sequence from phage KVP40 as a query. All proteins were searched using the ‘search’ function of MMseqs2 (release 12-113e3)^41^ using 2 iterations (parameter --num-iterations 2). Hits with an e-value lower than 10^−5^ were selected. Sequences from phage and prokaryotic genomes were separately filtered for redundancy by clustering sequences with identical length and at least 90% identity using the clust- hash option of MMseqs2^44^. Sequences shorter than 200 residues were discarded, and the remaining sequences were aligned using Clustal-Omega (v 1.2.4)^45^. Phylogenetic trees were built using IQtree (v. 1.6.5)^46^ with parameters -m LG -nt AUTO. Human Prps1 and Nampt were used as an outgroup for the Prs/Adps and Namat trees, respectively. Node support was computed using 1000 iterations of the ultrafast bootstrap function in IQtree (option -bb 1000)^23^. All trees were visualized with iTOL^47^. For the Prs/Adps tree, leaves were annotated based on the presence of a Namat homolog within ten genes upstream or downstream of Prs/Adps. For the Namat tree, each sequence was annotated based on the presence of a Prs/Adps or Nmnat homolog in the same phage genome.

## Acknowledgements

We thank members of the Sorek laboratory for comments on earlier versions of this manuscript. R.S. was supported, in part, by the European Research Council (grant no. ERC-AdG GA 101018520), Israel Science Foundation (MAPATS Grant 2720/22), the Deutsche Forschungsgemeinschaft (SPP 2330, Grant 464312965), a research grant from the Estate of Marjorie Plesset, the Ernest and Bonnie Beutler Research Program of Excellence in Genomic Medicine, Dr. Barry Sherman Institute for Medicinal Chemistry, Miel de Botton, the Andre Deloro Prize, and the Knell Family Center for Microbiology. I. O. was supported by Ministry of Absorption - New Immigrant program. E.Y. is supported by the Clore Scholars Program, and, in part, by the Israeli Council for Higher Education (CHE) via the Weizmann Data Science Research Center.

## Supplementary materials

### Supplementary Figures

### Supplementary Tables

**Supplementary Table 1**. Adps homologs

**Supplementary Table 2**. Namat homologs

**Supplementary Table 3**. Strains, plasmids and phages used in this study

**Supplementary Table 4**. Primers used in this study

**Supplementary Figure 1.**
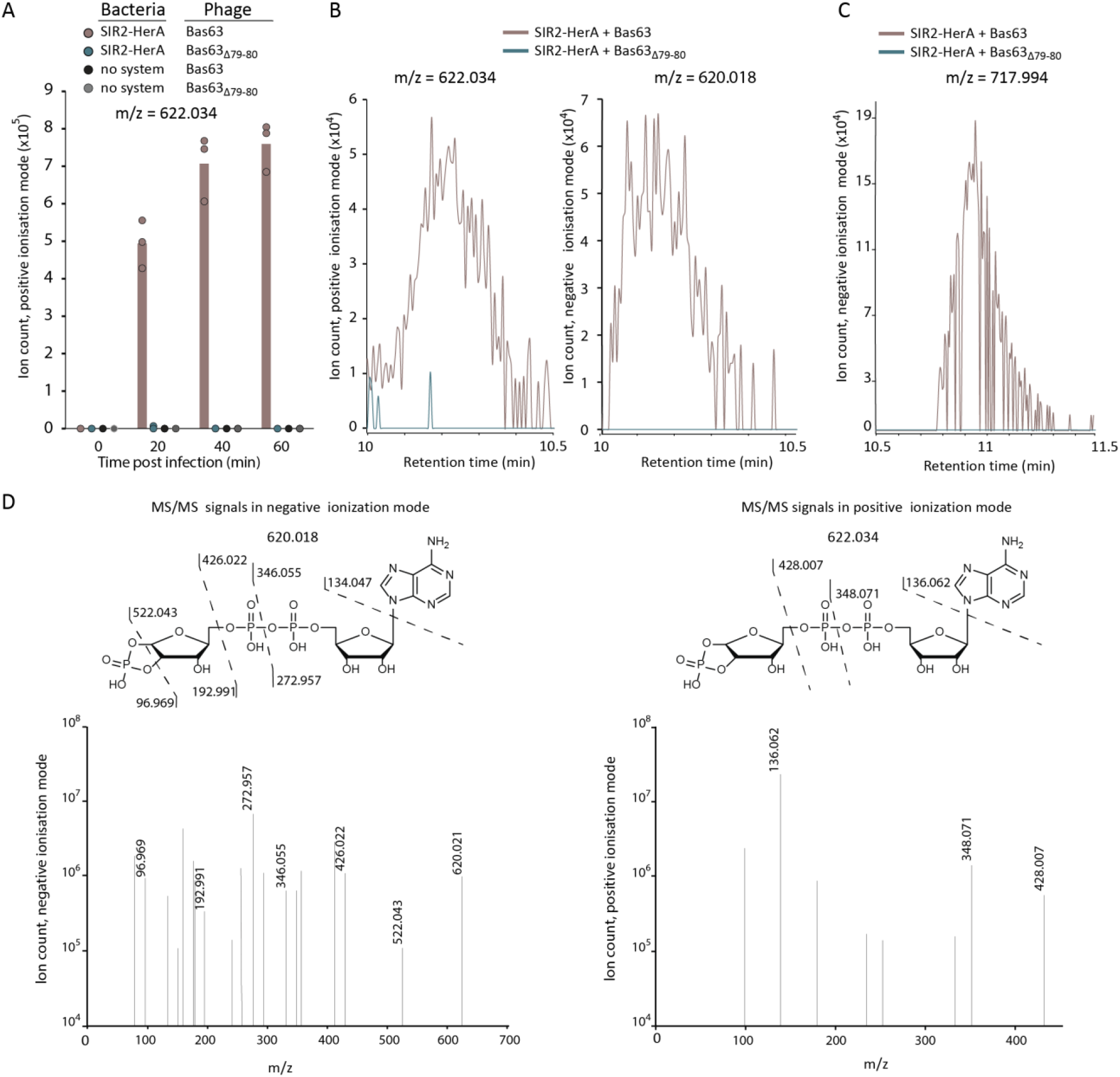
Mass spectrometry analysis of metabolites detected in lysates from infected cells. **A**. A unique molecule with an m/z value of 622.034 appears in SIR2-HerA cells infected by Bas63. Cells were infected at MOI=10. Bars represent the mean area under the curve (AUC) of three experiments, with individual data points overlaid. **B**. Extracted mass chromatograms of ions with an m/z value of 622.034 (positive ionization mode) and 620.021 (negative ionization mode) and retention time of 10.2 min. **C**. MS data in negative ionization mode for the same molecule presented in Figure 2B. **D**. MS/MS fragmentation spectra of the molecule with the m/z value 620.021 (negative ionization mode) and 622.034 (positive ionization mode). The hypothesized structure of the molecule and MS/MS fragments are presented.

**Supplementary Figure 2.**
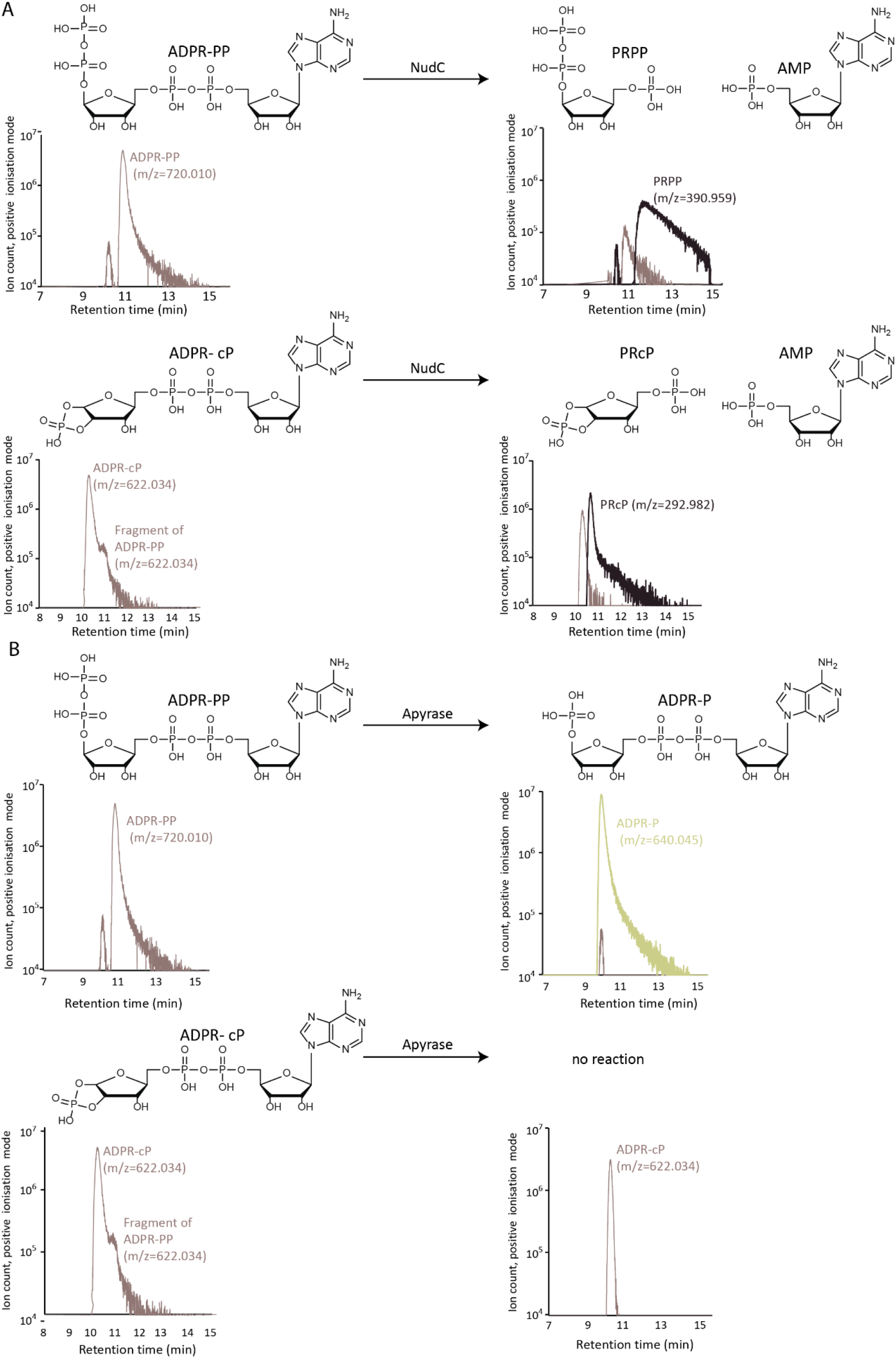
Enzymatic treatment of NARP1 products. A. Schematic of the reactions and mass- chromatograms of ADPR-PP and ADPR-cP following incubation with the enzyme NudC. Representative chromatograms of three replicates are presented. **B**. Schematic of the reactions and mass-chromatograms of ADPR-PP and ADPR-cP following incubation with the enzyme Apyrase. The peak with m/z 622.034 and retention time 11.0 is hypothesized to correspond to fragmentation of ADPR-PP by ionization in mass spectrometer. Representative chromatograms of three replicates are presented.

**Supplementary Figure 3.**
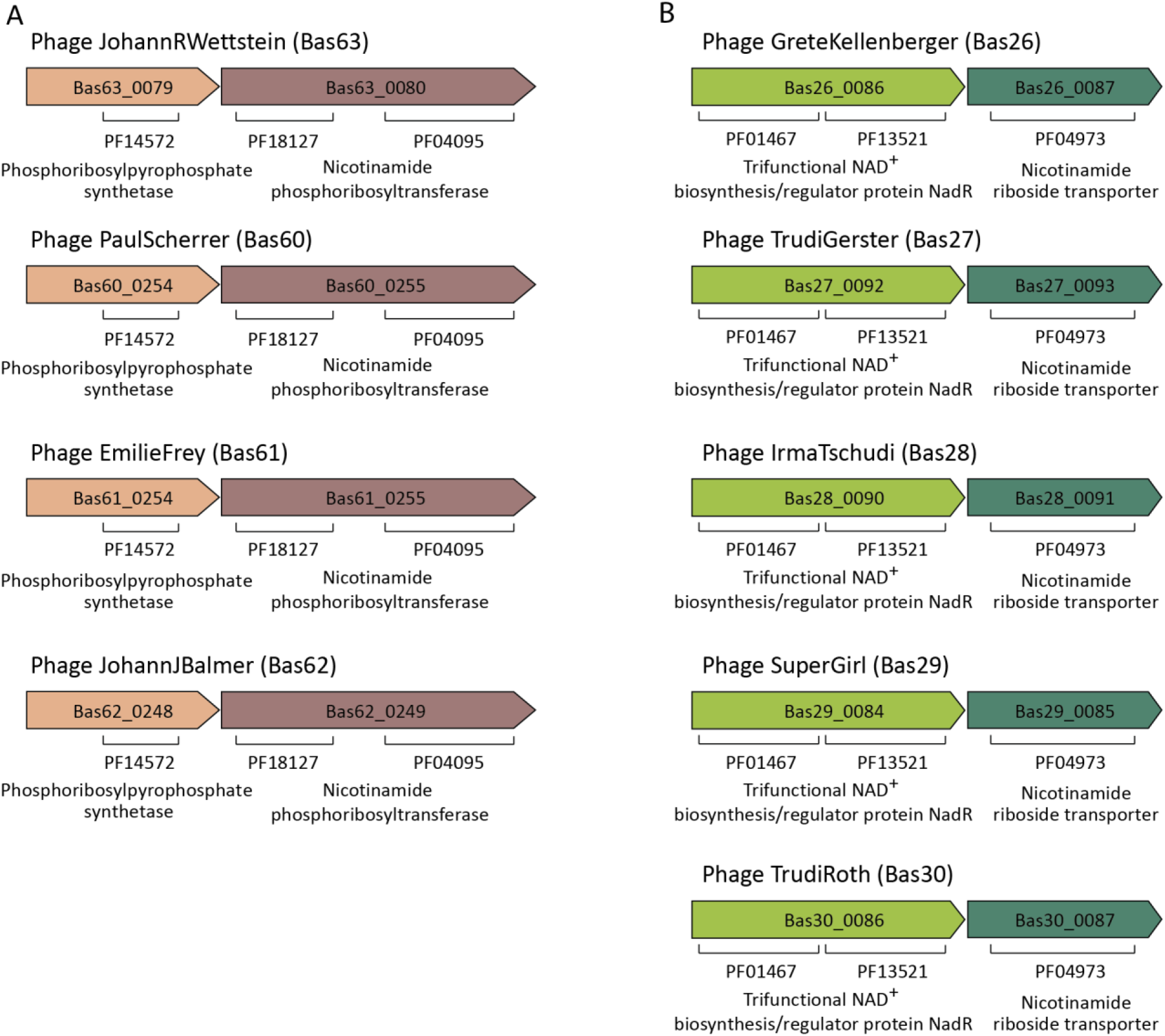
BASEL-collection phages encoding NAD reconstitution pathways. **A**. BASEL-collection phages that encode the NARP1 pathway. B. BASEL-collection phages that encode a two-gene operon predicted to comprise NadR and a transporter for nicotinamide riboside. This operon is hypothesized to comprise a phage NAD^+^ reconstitution pathway that was not examined in the current study.

